# An improved method to produce clinical scale natural killer cells from human pluripotent stem cells

**DOI:** 10.1101/614792

**Authors:** Huang Zhu, Dan S. Kaufman

**Author notes:** Corresponding author: Dan S. Kaufman, MD, PhD. University of California-San Diego, San Diego, California.

## Abstract

Human natural killer (NK) cell-based adoptive anti-cancer immunotherapy has gained intense interest with many clinical trials actively recruiting patients to treat a variety of both hematological malignancies and solid tumors. Most of these trials use primary NK cells isolated either from peripheral blood (PB-NK cells) or umbilical cord blood (UCB-NK cells), though these sources require NK cell collection for each patient leading to donor variability and heterogeneity in the NK cell populations. In contrast, NK cells derived human embryonic stem cells (hESC-NK cells) or induced pluripotent stem cells (hiPSC-NK cells) provide more homogeneous cell populations that can be grown at clinical scale, and genetically engineered if desired. These characteristics make hESC/iPSC-derived NK cells an ideal cell population for developing standardized, “off-the-shelf” immunotherapy products. Additionally, production of NK cells from undifferentiated human pluripotent stem cells enables studies to better define pathways that regulate human NK cell development and function. Our group previously established a stromal-free, two-stage culture system to derive NK cells from hESC/hiPSC in vitro followed by clinical-scale expansion of these cells using interleukin-21 expressing artificial antigen-presenting cells. However, prior to differentiation, this method requires single cell adaption of hESCs/hiPSCs which takes months. Recently we optimized this method by adapting the mouse embryonic fibroblast-dependent hESC/hiPSC to feeder-free culture conditions. These feeder-free hESC/hiPSCs are directly used to generate hemato-endothelial precursor cells. This new method produces mature, functional NK cells with higher efficiency to enable rapid production of an essentially unlimited number of homogenous NK cells that can be used for standardized, targeted immunotherapy for the treatment of refractory cancers and infectious diseases.

## Introduction

Human natural killer (NK) cells are an important part of innate immune system with ability to kill malignant and virally infected cells without MHC-restriction and without prior sensitization ^1^. NK cell cytotoxic activity against tumor cells or infected cells is mediated through a repertoire of germ-line encoded activating and inhibitory cell-surface receptors including killer-immunoglobulin receptors (KIRs), natural cytotoxicity receptors (NCRs) and the Fc Gamma receptor (FcγRIIIa) CD16a that mediates antibody-dependent cellular cytotoxicity (ADCC)^2, 3^. These important characteristics enable NK cells to function as allogeneic effector cells for treatment of refractory cancers and chronic infection diseases such as HIV.

Human embryonic stem cells (hESCs) and induced pluripotent stem cells (iPSCs) are ideal starting populations for the development of multiple cell lineages, including NK cells. hESC/iPSCs provide an model system to study the development NK cells in vitro^4, 5^. Previous studies from our group have shown that hESC/iPSC-derived NK cells have potent anti-tumor and anti-viral activities, providing a standardized cell-based treatment for these diseases ^6-8^.

Multiple clinical studies now demonstrate NK cells can effectively treat acute myeloid leukemia (AML) and other malignancies while not causing serious adverse effects such as graft-versus-host disease (GvHD) or cytokine release syndrome (CRS)^9, 10^. To date, most of NK cell-based adoptive immunotherapy clinical trials have used primary NK cells isolated from donor’s peripheral blood (PB-NK cells)^9, 10^, umbilical cord blood (UCB-NK cells)^11^ or the transformed NK cell line NK-92 ^12^. While each have demonstrate clinical efficacy, there are some shortcomings ^1^^3^. For example, PB-NK cells and UCB-NK cells are typically a heterogeneous mix of NK cells and other immune cells that can vary from donor to donor ^14^. NK-92 cells are aneuploid and need to be irradiated before use which limits the in vivo survival and expansion of these cells -- as is known to be a key determinant of anti-tumor activity^12^. In contrast, hESC-NK cells and hiPSC-NK cells are more homogenous and can generate essentially unlimited cells sources suitable for clinical use^8^. Importantly, hESC/hiPSC-NK cells exhibit similar phenotype, transcriptome and functions as primary NK cells ^6, 8^. Moreover, hESC/iPSC-NK cells can be routinely genetically modified by any of several methods including transposon and viral vectors^15^. Importantly, this genetic modification can be done on the undifferentiated pluripotent stem cells to produce a uniform population of NK cells with the desired effect. For example, the CRISPR/Cas9 or TALEN can be used to delete a gene of interest in hESCs or iPSCs to produce “knock-out” human NK cells. Key studies demonstrate the ability to utilize these genetic reporter systems expressed in hESCs and iPSCs to track and regulate development of specific blood cell lineages ^4, 5, 16^.

The methods to generate NK cells from human pluripotent stem cells has evolved in the last decade ^6, 17, 18^. Our initial studies used a stromal cell dependent two-stage culture system^6^. Briefly, to obtain CD34^+^CD45^+^ hematopoietic progenitor cells, undifferentiated hESCs were cultured on stromal cell line (e.g. S17 or M2–10B4 cells) using media containing fetal bovine serum (FBS). Then hematopoietic progenitor cells were sorted and moved to a second stromal cell line (e.g. AFT024 or EL08-1D2 cells) in media supplemented with SCF, Fms-like tyrosine kinase 3 ligand (FLT3L), IL-3, IL-15, IL-7 to direct differentiation towards NK cells. Later, our group optimized the protocol by adapting a “spin-embryoid body (EB)” method to derive hematopoietic progenitor cells in defined serum-free, stromal-free conditions^8, 18^. After 11 days culture, hematopoietic progenitor cell-containing EBs were transferred to feeder-free plates in NK differentiation media containing SCF, FLT3L, IL-3, IL-15 and IL-7 for 4 weeks to generate CD45^+^CD56^+^ NK cells. hESC-NK/iPSC-NK cells obtained using these methods express activating and inhibitory receptors similar to PB-NK cells^8^. More importantly, these NK cells exhibit potent anti-tumor and anti-viral activity both in vitro and in vivo. These functions include effective elimination of tumor cells xenografted in immune-deficient mice and inhibition of HIV-infected targets in an in vivo SCID-hu model ^6, 7, 15, 19^. hESC-NK/iPSC-NK cells can be further expanded to clinical scale by using irradiated human stimulator cells, as previously utilized to effectively expand PB-NK or UCB-NK cells ^20^. Notably, this method requires an essential single cell adaptation step to enable hESC/iPSC survival as single cells ^18, 21^. This process is not uniform and typically requires 12-15 single cell passages that take several weeks.

Here, we describe an improved method for the derivation of NK cells from human pluripotent stem cells with higher efficiency and less time than existing protocols ^18^. NK cells derived using this method have similar growth rate, phenotype and activity compared to PB-NK cells and UCB-NK cells generated by existing methods. In summary, this improved method allows efficient, rapid and reproducible production of hESC-/iPSC-NK cells from different pluripotent stem cell lines without the requirement of time-consuming single cell adaption process to facilitate clinical scale production and translation of hESC/iPSC-NK cells.

Figure 1 provides a schematic diagram to compare the old and new methods. Prior to the generation of EBs, MEF-dependent undifferentiated hESC/iPSC are passaged on Matrigel-coated plates in mTeSR media (feeder free conditions) for 1 week, though this step is not needed for hESC/iPSC already maintained in feeder-free conditions. The feeder-free hESC/iPSC can be directly used to form EB in the presence of Rho-associated protein kinase inhibitor (ROCKi) which decreases the cellular stress response and the apoptotic cell death in stem cell cultures ^22^. Indeed, these new conditions produce more hESC/iPSC-derived CD34^+^ hematopoietic progenitor cells in only 6 days (Figure 2 G). Following the generation of the hemato-endothelial precursor cells in the EBs, NK cell differentiation is supported using a stromal cell-free protocol similar to the existing protocol ^18^.

**Figure 1.**
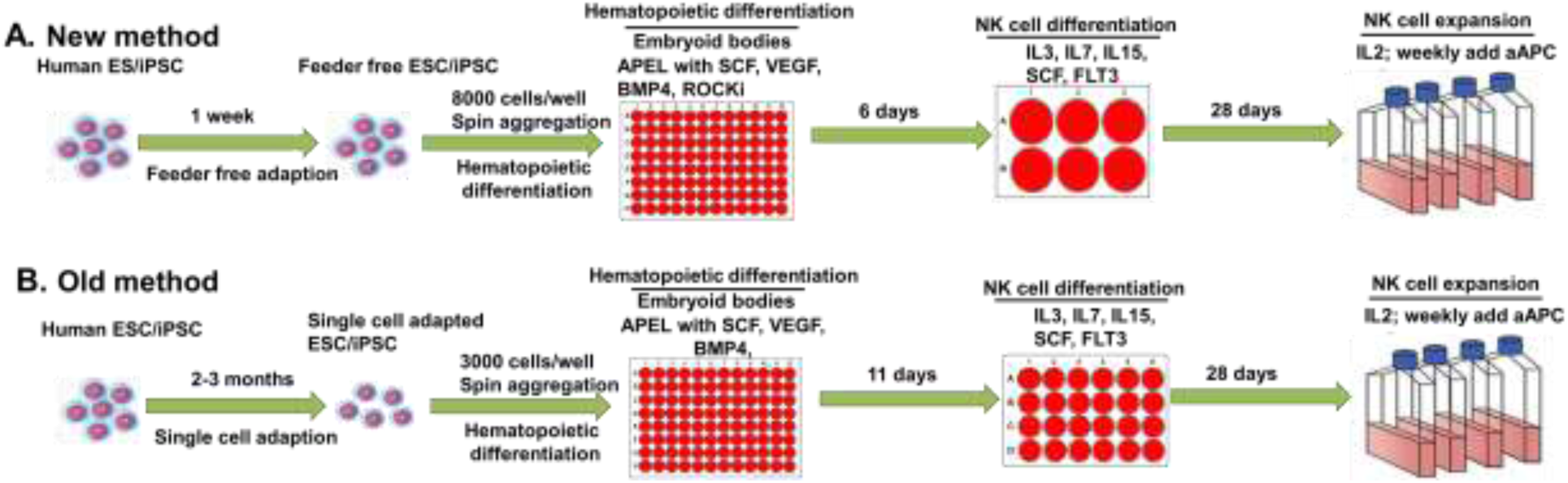
Schematic diagram of new method. (A.) and old method ^**18**^ (B.) for in vitro NK cell differentiation from human embryonic stem cells (ESC) or induced pluripotent stem cells (iPSC). Briefly, the old method requires a single cell adaption process for ESC/iPSC which takes 2-3 months. While the new method starts with feeder free adapted ESC/iPSC and use ROCKi to help EB formation. Both methods use spin EBs (11 days for old method and 6 days for the new method) to generate hematopoietic progenitor cells (CD34+ cells). Then EBs are directly transferred into NK cell differentiation conditions. Mature and functional NK cells will develop after 3-5 weeks and can be expanded using IL2 and aAPCs.

**Figure 2.**
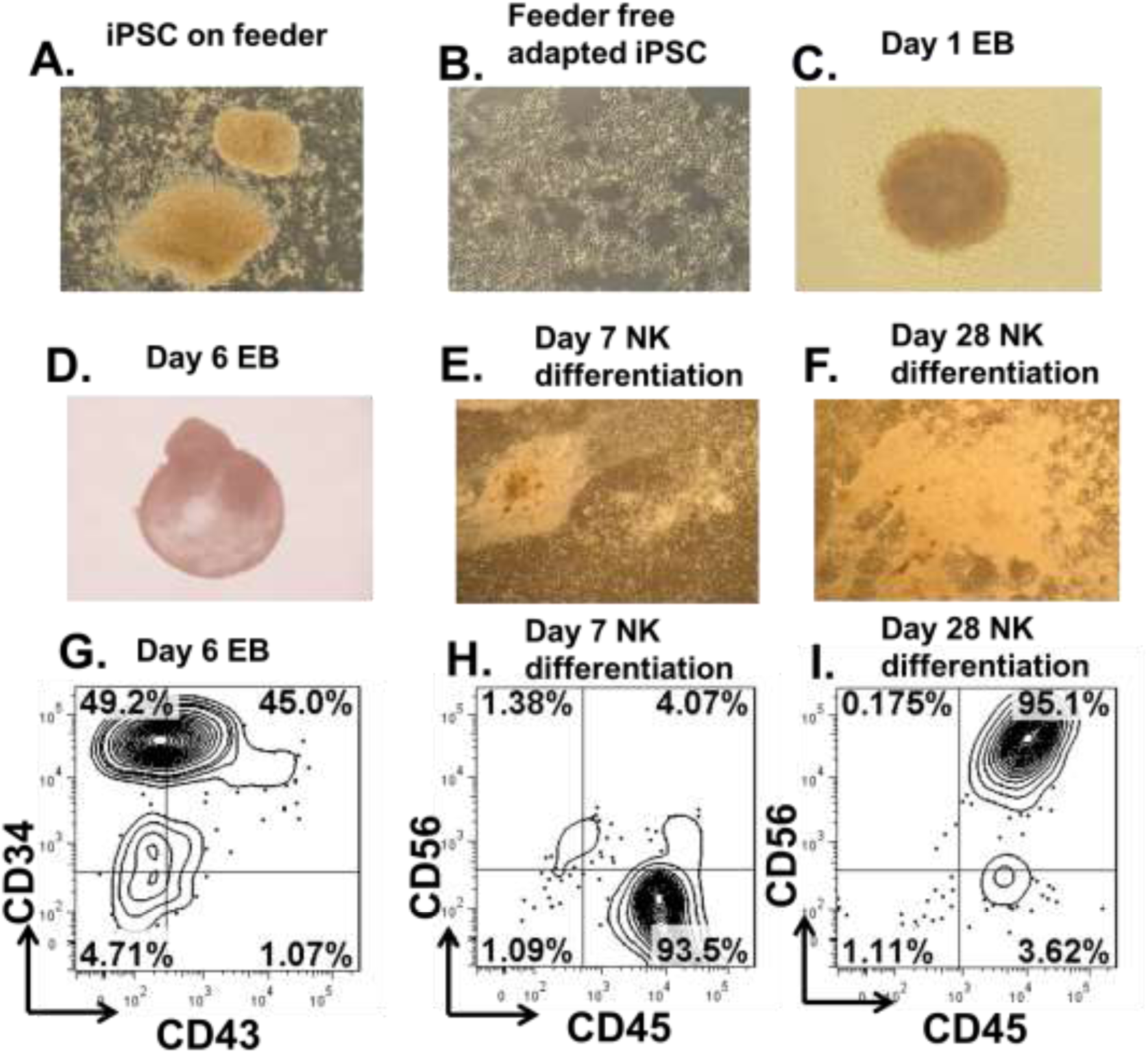
Phase microscope images and phenotypes of differentiating NK cells from human pluripotent stem cells at different stages. **A**= Undifferentiated iPSCs on MEFs. **B**= Feeder-free undifferenatiated iPSCs. **C**= Day 1 Embryoid body. **D and G**= Day 6 embryoid body. **E and H**= Day 7 NK cell differentiation. **F and I=** Day 28 NK cell differentiation. Original magnification × 100 for A-D, and ×20 for E, F.

## Materials

### Cell lines

We have successfully produced NK cells from several different iPSC lines, including those derived from human fibroblasts (FiPSC), human peripheral blood mononuclear cells (PBiPSCs)^23^, or human CD34+ cells isolated from umbilical cord blood (UiPSC)^8^. Human ES cell line H9 was obtained from Wicell, Madison WI. K562 aAPC cells expressing membrane-bound IL-21 (Clone 9. mbIL-21) ^20^ were kindly provided by Dr. Dean Lee and Dr. Lawrence Cooper, MD Anderson Cancer Center, Houston, TX.

### Cell culture media and reagents

#### Undifferentiated hESC and hiPSCs

culture in mTsER™1 (STEMCELL Technologies, Catalog # 85850).

#### K562 aAPC cells and Moml13 cells

Culture in RPMI-1640 (ThermoFisher Scientific, Catalog #11875-085), 2 mM L-glutamine (ThermoFisher Scientific, Catalog #25030081), 10 % FBS, and 1 % penicillin/streptomycin (ThermoFisher Scientific,Catalog # 15140122); Spin EB differentiation medium: STEMdiff™ APEL™2 Medium (STEMCELL Technologies,Catalog # 05270).

#### NK cell differentiation medium^18^

56.6 % DMEM+ GlutaMAX™-I (Life Technologies, Catalog # 10566-016), % F12+ GlutaMAX™-I (Life Technologies, Catalog # 31765035), 15 % heat-inactivated human AB serum (Valley Biomedical, Catalog # HP1022 HI),1 % P/S, 2 mM L-glutamine, 25 uM β-mercaptoethanol (Sigma, Catalog # M6250), 5 ng/mL sodium selenite (Sigma-Aldrich, Catalog # S5261), 50 uM ethanolamine (MP Biomedicals, Catalog #194658),20 mg/L ascorbic acid (Sigma-Aldrich, Catalog # A-5960), 5 ng/mL IL-3 (PeproTech, Catalog # 200-03), 20 ng/mL SCF, 20 ng/mL IL-7 (PeproTech, Catalog # 200-07), 10 ng/mL IL-15 (PeproTech, Catalog # 200-15), and 10 ng/mL Flt3 ligand (Flt3L) (PeproTech, Catalog #300-19), Store at 4 °C in the dark;

#### NK cell expansion medium

RPMI-1640, 10 % FBS, 2 mM L-glutamine, 1 % P/S, and 50 U/mL IL-2 (PeproTech, Catalog # 200-02).

#### Antibodies^8^

NKp46-PE, NKp44-PE, CD56-APC, NKG2D-PE, TRAIL-PE, FAS ligand-PE are from Becton, Dickinson and Company (Franklin Lakes, NJ, http://www.bd.com). CD16-APC is obtained from eBioscience Inc. (San Diego, CA, http://www.ebioscience.com). CD158a/h-PE, CD158j-PE, CD158i-PE, CD158e1/e2, and CD159a-PE are from Beckman Coulter (Fullerton, CA, http://www.beckmancoulter.com).

#### Other material

Matrigel Matrix (Corning, Catalog #354246); TrypLE Select (Life Technologies, Catalog # 12563011); Collagenase Type IV (STEMCELL Technologies, Catalog #07909); ROCKi: Y-27632 (Sigma, Catalog # SCM075); 96-well round bottom plates (For EB formation; NUNC, cat. no. 262162); 24-well or 6-well tissue culture plates (Corning, Catalog # 3527/3506); CellEvent® Caspase-3/7 Green Detection Reagent (Thermal Fisher Scientific, C10423); CellTrace™ Violet (Thermo-Fisher Scientific, C34557); S T X™ AADvanced™ dead cell stain solution (Thermal Fisher Scientific, S10349);

## Methods

### Feeder free adaptation of feeder dependent hESC/iPSC

Before generating EBs, undifferentiated feeder-dependent hESC/iPSC need to be transferred to feeder-free conditions with the use of mTeSR™1 on Matrigel (Corning) coated plates, as previously described ^24^. If hESC/iPSCs are already maintained on feeder-free conditions, skip this step. It is critical to make sure the hESC/iPSC are not differentiated before and after adaptation. This step will change the hESC/iPSC from clump culture to monolayer culture and increase efficiency of making single cell suspension. Cells should be able to adapt to feeder-free culture within 1 - 2 passages and then exhibit morphology consistent with feeder-free human pluripotent stem cells (Figure 2. A and B). The following instructions are used for passaging cells from one well of a 6-well plate.

1. Before passaging, coat new plate with 1× Matrigel (see instruction below) and incubate in 37°C incubator for 2-4 hours before use. Preparing 1× Matrigel:
  1.1 Thaw 5 mL stock vial of Matrigel at 4°C overnight.
  1.2 Quickly aliquot Matrigel using pre-chilled tips into pre-chilled Eppendorf tubes.
  1.3 Store at −80°C for up to 6 months.
  1.4 To prepare 1× Matrigel working solution, re-suspend one Matrigel aliquot at 1:100 ratio into cold DMEM/F12. 1× Matrigel working solution can be stored at −4°C for up to 2 weeks.
  1.5 Coat plate with sufficient volume to cover surface (e.g. use 1mL per well of 6 well plate).
  1.6 Incubate in 37°C incubator for 2 to 4 hours before use.
2. After making sure the feeder dependent hESC/iPSC is free of differentiated colonies and at right confluence (70-80%), aspirate media and pass cells by incubating cells for 15 min at 37 °C in pre-warmed Collagenase Type IV (1 mg/mL).
3. Aspirate the collagenase and wash cells once with 3 ml PBS. Then add 1 mL of mTeSR™1 medium. Then gently detach the colonies by scraping with a serological glass pipette.
4. Collect detached aggregates and add 3 ml of mTeSR™1 medium. To break up the cell aggregates to appropriate size (50-200uM), gently pipette up and down 2-3 times using a 5 mL serological pipette.
5. Plate cell aggregate mixture 1:2 onto Matrigel pre-coated plate.
6. Change media daily and repeat procedure once when cells become 70-80% confluent. Most of the feeder cells should be eliminated after 2 passages.
7. After 2 passages with collagenase, passage cells by incubating cells for 5 min at 37°C in pre-warmed TrypLE Select.
8. Carefully dissociate cells aggregates to single cell using 1 mL micropipette and transfer cells to a 15 mL conical tube. Add additional 5 ml medium (DMEM/F12 + 10%FBS).
9. Pass cell mixture through a 70 uM cell strainer to remove aggregates.
10. Centrifuge cells at 300 × *g* for 5 minutes.
11. Aspire supernatant and wash cells once with 5 ml PBS.
12. Re-suspend cells with mTeSR™1 medium and plate cells 1 2 onto Matrigel pre-coated plate. Cells can be used for spin EB formation when become confluent.

### Generation of Hematopoietic Progenitor Cells from hES/iPS Cells by Spin EB Formation

Our group adapted a ‘spin EB’ protocol to generate hematopoietic progenitors ^8, 21, 25^. Cells inside EBs can form endothelial and mesenchymal stromal cells to support following lymphocyte development, eliminating the use of xenogeneic stromal cells such as OP9 ^18^. While there is variability between different hESC and iPSC lines, we can consistently obtain >50% CD34+ cells for most of lines using this method (Figure 2) (Note 1). The following instructions are used for collecting cells from one well of a 6-well plate for EB formation.

1. 2 days before spin EB formation, pass 200,000 feeder free hESC/iPSCs onto 1 well of Matrigel coated 6-well plate which should reach 70–80 % confluence on the day of spin EBs set up. Generally 1 well of a 6-well plate are sufficient for 2 plates of spin EB.
2. Aspirate culture medium and detach cells by incubating with 1 ml incubating cells for 5 min at 37°C in pre-warmed TrypLE Select.
3. Carefully dissociate cells aggregates to single cell using 1 mL micropipette and transfer cells to a 15 mL conical tube. Add additional 5 ml medium (DMEM/F12 + 10%FBS). Pass cell mixture through a 70 uM cell strainer to remove aggregates.
4. Centrifuge cells, remove supernatant, wash cells once with 5 ml PBS. Then resuspend the cells in 1 ml APEL media.
5. Count cells and dilute cells to appropriate density using APEL media containing cytokines plus 10 uM ROCKi (Y-27632) to be used for plating. Typically, cells are seeded at 8000 cells/well in 100 ul media (80,000 cells/ml) in the 96-well plates (Note 2).
6. Pipet 100 µl of the cell suspension into 96-well plates using a multi-channel pipet. Centrifuge 96-well plates at 300 × g for 5 minutes, and incubate the plates at 37 °C, 5 % CO2 for 6 days. We have found 6 days are enough to give rise to >50% CD34+ cells for most of hESC and iPSC lines.

### Derivation of human NK cells from spin EB

Spin EBs can be transferred into either 24-well plates or 6-well plates coated with 2% gelatin or without coating. We have found that 6-well plate is more suitable for medium change and 2% gelatin can help EBs attachment.

1. Before transferring EBs, coat new 6-well plates with 2% gelatin (see instruction below) and incubate in 37°C incubator for 2-4 hours before use. **Preparing 2% gelatin coated plate:**
  1.1 Prepare a 2% (w/v) solution by dissolving gelatin (SIGMA #G1890) in tissue culture grade water
  1.2 Sterilize by autoclaving at 121°C, 15 psi for 30 minutes. Let cool.
  1.3 Coat culture surface with 5-10 µL gelatin solution/cm2 (about 1 ml per well of 6-well plate).
  1.4 Allow to incubate 37°C for one hour, then aspirate remaining gelatin solution before introducing cells and medium
2. Add 3 ml NK differentiation medium (as above) containing all of the cytokines to each well of gelatin-coated 6-well plate.
3. Spin EBs are directly transferred into 6-well plates on day 6. Carefully transfer EBs from 96-well plates to a 10 cm dish using 10 ml serological pipette and remove most of medium (Note 3). Then add 2 ml of NK differentiation medium containing all of the cytokines. Distribute 14-16 EBs into each well of the 6-well plate using 1 mL micropipette. Typically, EBs from 1 96-well plate can make 1 6-well NK differentiation plate.
4. Medium changes are done every 5–7 days (Note 4). Following the first week of the NK cell differentiation, IL-3 is no longer added to the media.
5. Continue medium changes for 3-4 weeks and check expression of NK cell makers (CD45+CD56+) by flow cytometry on suspension cells (Figure 2 E-I). Cells can be collected passing them through a 70-µm filter to remove any clumps when >80% of suspension cells are CD45+CD56+ (Note 5).

### Expansion of hESC-/iPSC-Derived NK Cells to clinical scale

Typically, we can obtain 2-20 × 10^6^ NK cells from 1 6-well plate. To further expand the NK cells for downstream applications, artificial antigen-presenting cells (aAPCs) are used to generate >10^9^ NK cells (Figure 3).

**Figure 3.**
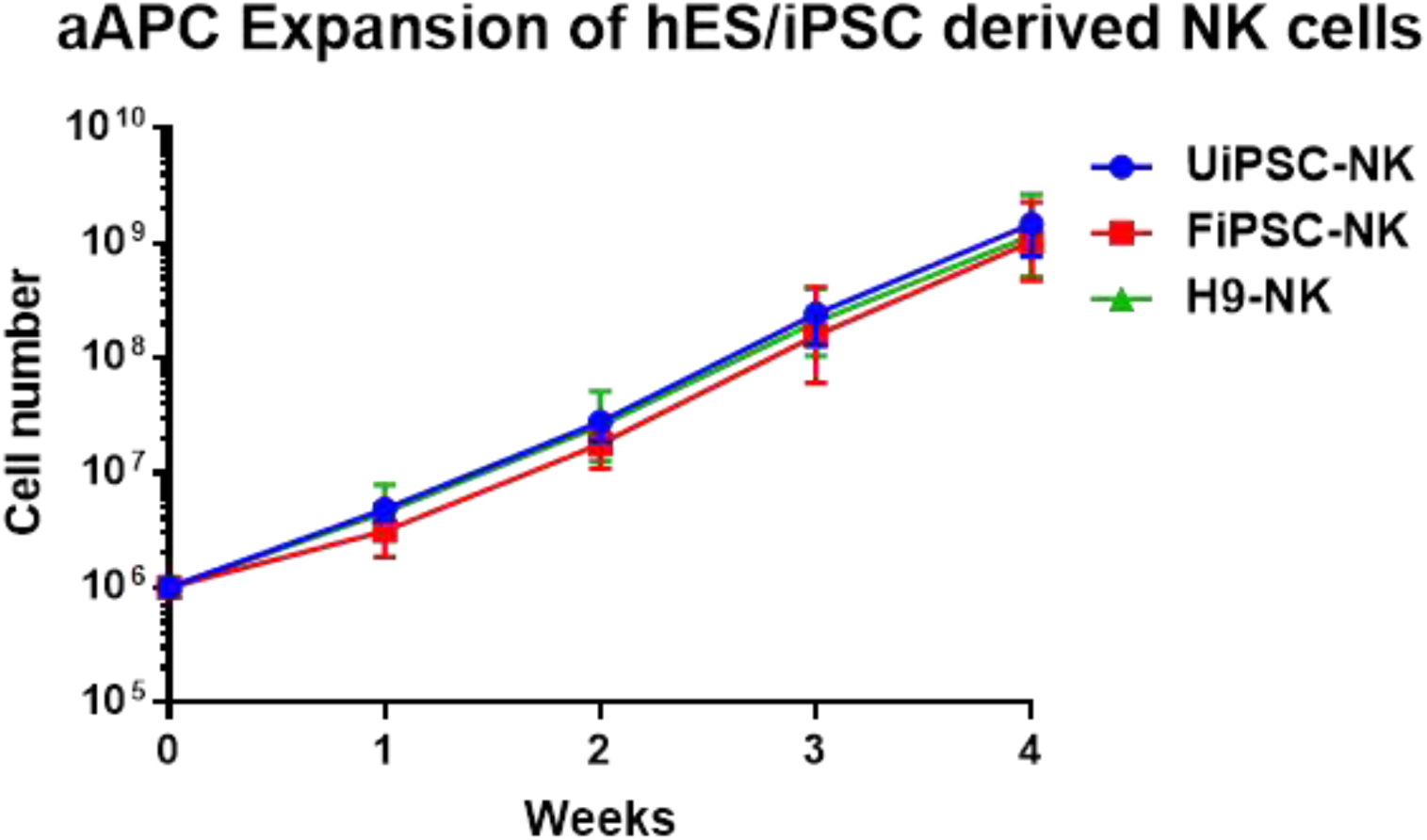
Growth curve of NK cells derived from hES/iPSC using method presented in this chapter. 2 iPSC lines (iPSC reprogrammed from umbilical CD34+ cells (UiPSC) and from fibroblast (FiPSC)) and 1 hESC line H9-derived NK cells were transferred from NK differentiation conditions and placed in aAPC for expansion. Cells were stimulated with aAPCs weekly and cell number reached >10^9^ from 10^6^ starting cell number in 4 weeks. Each line represents mean of 3 independent experiments.

1. We use membrane-bound IL-21 expressing (mbIL-21) K562 cells^8, 20^ as aAPCs to stimulate NK cell expansion. Before adding to NK cells, aAPCs are irradiated with 10,000 rads and made as frozen stocks.
2. After collecting from 6-well plate, NK cells are maintained in NK expansion medium containing 50 U/mL IL-2 (add freshly) at a density of 3 × 10^5^ cells/mL. Irradiated aAPCs are thawed and added to NK cells at a ratio of 1:1. Media are changed every 3–4 days containing 50 U/mL of freshly added IL-2.
3. hESC-derived or iPSCs derived NK cells can be expanded for >3 months without a decrease in cell viability or cytolytic activity.

### Phenotypic and functional characterization of hESC/iPSC-derived NK cells

hESC/iPSC-derived NK cells can be phenotyped for surface antigens including CD56, CD45, CD16, NKG2D, NKp44, NKp46, TRAIL, FasL etc. We normally characterize phenotypes and killing activity of hESC/iPSC-derived NK cells after 3 weeks of expansion. Functions of NK cells can be accessed in vitro by measurement of direct cytolytic activity tumor cells (such as K562) by, ^51^Cr-release, Caspase3/7 flow cytometric assay or immunological assays for cytotoxic granule or cytokine release. If desired, anti-tumor activity can be assessed in vivo using xenograft models ^6, 19^. NK cells developed using this method show a mature NK cell phenotype and cytotoxicity (Figure 4).

**Figure 4.**
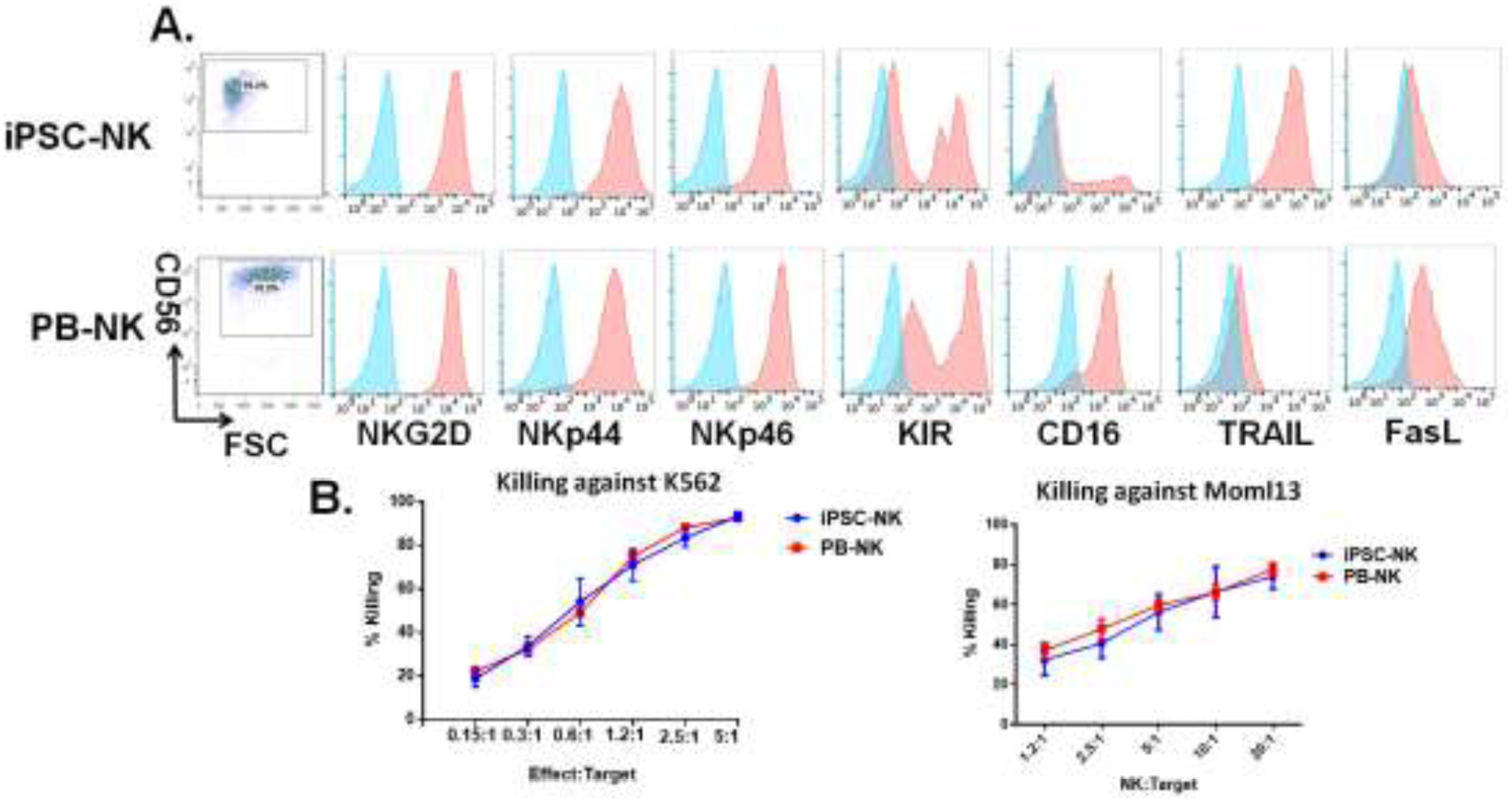
iPSC-NK cells are phenotypically mature and have similar killing activities against tumor targets as PB-NK cells. **A.** iPSC-NK cells and PB-NK cells were stained for a panel of NK cell receptors. Expression of each marker is shown by representative flow cytometry plots. **B.** In vitro killing against K562 cells and M LM13 cells was determined using CellEvent™ Caspase-3/7 Green Flow Cytometry Assay. Different amount of target cells were incubated with either iPSC-NK cells or PB-NK cells (effector:target ratio from 0.15:1 to 5:1 for K562, 1.2:1 to 20:1 for MOLM13 cells) for 4 hours. Data are represented as mean ± SEM.

1. We typically test the expression of following receptors by flow cytometry: CD56, CD16, CD94, NKG2D, NKp44, NKp46, KIR, FasL and TRAIL.
2. After analyzing phenotypes, functional of hESC/iPSC-derived NK cells are evaluated using flow cytometry based Caspase 3/7 assay against tumor targets:
  2.1 Target cells were counted and pre-stained with CellTrace™ Violet at a final concentration of 5µM in PBS for 15min at 37°C.
  2.2 After staining, target cells were washed in complete culture medium prior to being mixed with NK cell cultures at the indicated effector to target (E:T) ratios.
  2.3 Incubate NK cells and target cells mix at 37 degree for 3.5 hrs.
  2.4 Add CellEvent® Caspase-3/7 Green Detection Reagent for an additional 30 minutes of culture for a total incubation time of 4 hours.
  2.5 During the final 5 minutes of staining, add 1 µL of S T X™ AADvanced™ dead cell stain solution and mixed gently
  2.6 Cells were then analyzed by flow cytometry

## Notes

### Note 1

We are able to obtain >50% CD34+ cells for most of iPSC and ES lines tested using this method. We also found that EBs containing >10% CD34+ cells can be successfully differentiated to mature NK cells.

### Note 2

8000 cells/well works for most of cell lines. But the cell number can be optimized by individual investigators using different pluripotent stem cell lines depending on the growth rate of the cell line. We seed more cells than our previous protocol ^18^ (3000 cells/well) as we reduced the EB formation time from 11 days to 6 days.

### Note 3

It is critical to remove most of the medium and re-suspend with NK cell differentiation medium.

### Note 4

After 14 days, medium needs to be changed every 3 days.

### Note 5

It usually takes 4 weeks to obtain >90% CD45+CD56+ cells from spin EBs NK cells for most of ES or iPSC lines (Figure 2). Some cell line might only require 3 weeks and some line may take longer (about 5 weeks).

## Result

### Single cell adaption of ESC/iPSC is not required for generation of hematopoietic progenitor cells and NK cells

Our group adapted a ‘spin EB’ protocol to generate hematopoietic progenitors ^8, 21, 25^. Cells inside EBs can form self-stromal cells to support following lymphocyte development, eliminating the use of xeno-derived stromal cells such as OP9 ^18^. However, this method requires a single cell adaption process for ESC/iPSC which takes 2-3 months. To test whether the single cell adaption of ESC/iPSC is essential for generation of hematopoietic progenitor cells, we tried EB formation from human iPSC without single cell adaption (Figure 1). First, we culture the iPSCs without feeders for 1 week (Figure 2 A,B). ROCKi was added to prevent cell death and help EB formation. Intact EBs can form from hiPSCs without single cell adaption (Figure 2 C, D). Notably, most of cells in the EBs (day6) are CD34^+^ and about half of them are CD34^+^CD45^+^ (Figure 2 G). We then transferred day 6 EB to NK cell differentiation medium (see method section). After 7 days in NK cell differentiation medium, we found lots of round cells (suspension) around the EBs (Figure 2 E). Most of the suspension cells were CD45^+^CD56^-^ at this time point (Figure 2 H). And most of the suspension cells become CD56+ after 28 days in NK cell differentiation medium (Figure 2 F,I). Together, these results demonstrated that Single cell adaption of ESC/iPSC is not required for generation of hematopoietic progenitor cells and NK cells.

### Human ESC/iPSC derived NK cells using the new method are phenotypically mature and functional

We then generated NK cells from 2 iPSC lines (iPSC reprogrammed from umbilical CD34+ cells (UiPSC) and from fibroblast (FiPSC)) and 1 hESC line H9-derived NK cells using the new method described in Figure 1. All the NK cells show similar growth rate when stimulating with aAPC weekly (Figure 3). Importantly, hiPSC-derived NK cells using this method express NK cells receptors including NKG2D, NKp44, NKp46, TRAIL, FasL, similar to that of primary NK cells (Figure 4A). About 20% iPSC-NK cells were CD16+ which was consistent with previous publications. Moreover, hiPSC-derived NK cells show similar killing against tumor cells such as K562 and Molm13 (Figure 4B). These results demonstrated that Human ESC/iPSC derived NK cells using the new method are phenotypically mature and functional similar to primary NK cells.

## Discussion

Here we optimized the method to derive NK cells from human ESC/iPSC. The improved method does not require the need to adapt ESC/iPSC to single cells, a process that takes 2-3 months. This new method takes about 5 weeks to generate large numbers mature NK cells from ESC/iPSC. Notably, hESC/iPSC-derived NK cells using this method can be efficiently expanded to clinical scale. Importantly, these cells are phenotypically mature and functional similar to primary NK cells.

ROCK inhibition was reported to improve the cloning efficiency and survival upon dissociation of hES cells without altering their karyotype or pluripotency^26^. Here we used Y-27632, a specific inhibitor of Rho kinase (ROCK) activity, to help EB formation. Without Y-27632, most human ESC/iPSC without single cell adaption can’t form intact EBs. In this new protocol, we transferred EB to NK cell differentiation medium at day 6 as most of cells in the EBs were CD34^+^ at this time point. Although most of cells were still CD45^-^, they become CD45^+^ shortly after culturing in NK cell differentiation medium.

Cell-based therapies for treatment of relapsed/refractory cancers have been gaining in interest and importance. In addition to studies using CAR-T cells, clinical trials using NK cells isolated from peripheral blood or umbilical cord blood have been rapidly expanding^13^. However, these sources of NK cells are difficult to engineer and vary from donor to donor. iPSC-derived NK cells are being translated to clinical trials to now provide a standardized, “off-the-shelf” therapy that can be routinely engineered to target and better treat relapsed/refractory malignancies^13^. The improved method allows efficient, rapid and reproducible production of hESC-/iPSC-NK cells from different pluripotent stem cell lines without the requirement of time-consuming single cell adaption process thus facilitate clinical scale production and translation of hESC/iPSC-NK cells.

## References

1. Vivier, E., Raulet, D. H., Moretta, A., Caligiuri, M. A., Zitvogel, L., Lanier, L. L., Yokoyama, W. M. & Ugolini, S. (2011). Innate or adaptive immunity? The example of natural killer cells. Science 331, 2011.

2. Morvan, M. G. & Lanier, L. L. (2016). NK cells and cancer: you can teach innate cells new tricks. Nat Rev Cancer 16, 2016.

3. Jing, Y., Ni, Z., Wu, J., Higgins, L., Markowski, T. W., Kaufman, D. S. & Walcheck, B. (2015). Identification of an ADAM17 cleavage region in human CD16 (FcgammaRIII) and the engineering of a non-cleavable version of the receptor in NK cells. PLoS One 10, 2015.

4. Angelos, M. G., Ruh, P. N., Webber, B. R., Blum, R. H., Ryan, C. D., Bendzick, L., Shim, S., Yingst, A. M., Tufa, D. M., Verneris, M. R. & Kaufman, D. S. (2017). Aryl hydrocarbon receptor inhibition promotes hematolymphoid development from human pluripotent stem cells. Blood 129, 2017.

5. Ferrell, P. I., Xi, J., Ma, C., Adlakha, M. & Kaufman, D. S. (2015). The RUNX1 +24 enhancer and P1 promoter identify a unique subpopulation of hematopoietic progenitor cells derived from human pluripotent stem cells. Stem cells 33, 2015.

6. Woll, P. S., Grzywacz, B., Tian, X., Marcus, R. K., Knorr, D. A., Verneris, M. R. & Kaufman, D. S. (2009). Human embryonic stem cells differentiate into a homogeneous population of natural killer cells with potent in vivo antitumor activity. Blood 113, 2009.

7. Ni, Z. Y., Knorr, D. A., Clouser, C. L., Hexum, M. K., Southern, P., Mansky, L. M., Park, I. H. & Kaufman, D. S. (2011). Human Pluripotent Stem Cells Produce Natural Killer Cells That Mediate Anti-HIV-1 Activity by Utilizing Diverse Cellular Mechanisms. J Virol 85, 2011.

8. Knorr, D. A., Ni, Z., Hermanson, D., Hexum, M. K., Bendzick, L., Cooper, L. J., Lee, D. A. & Kaufman, D. S. (2013). Clinical-scale derivation of natural killer cells from human pluripotent stem cells for cancer therapy. Stem Cells Transl Med 2, 2013.

9. Miller, J. S., Soignier, Y., Panoskaltsis-Mortari, A., McNearney, S. A., Yun, G. H., Fautsch, S. K., McKenna, D., Le, C., Defor, T. E., Burns, L. J., Orchard, P. J., Blazar, B. R., Wagner, J. E., Slungaard, A., Weisdorf, D. J., Okazaki, I. J. & McGlave, P. B. (2005). Successful adoptive transfer and in vivo expansion of human haploidentical NK cells in patients with cancer. Blood 105, 2005.

10. Romee, R., Rosario, M., Berrien-Elliott, M. M., Wagner, J. A., Jewell, B. A., Schappe, T., Leong, J. W., Abdel-Latif, S., Schneider, S. E., Willey, S., Neal, C. C., Yu, L., Oh, S. T., Lee, Y. S., Mulder, A., Claas, F., Cooper, M. A. & Fehniger, T. A. (2016). Cytokine-induced memory-like natural killer cells exhibit enhanced responses against myeloid leukemia. Sci Transl Med 8, 2016.

11. Verneris, M. R. & Miller, J. S. (2009). The phenotypic and functional characteristics of umbilical cord blood and peripheral blood natural killer cells. British journal of haematology 147, 2009.

12. Tonn, T., Schwabe, D., Klingemann, H. G., Becker, S., Esser, R., Koehl, U., Suttorp, M., Seifried, E., Ottmann, O. G. & Bug, G. (2013). Treatment of patients with advanced cancer with the natural killer cell line NK-92. Cytotherapy 15, 2013.

13. Zhu, H., Lai, Y. S., Li, Y., Blum, R. H. & Kaufman, D. S. (2018). Concise Review: Human Pluripotent Stem Cells to Produce Cell-Based Cancer Immunotherapy. Stem Cells 36, 2018.

14. Passweg, J. R., Tichelli, A., Meyer-Monard, S., Heim, D., Stern, M., Kuhne, T., Favre, G. & Gratwohl, A. (2004). Purified donor NK-lymphocyte infusion to consolidate engraftment after haploidentical stem cell transplantation. Leukemia 18, 2004.

15. Ni, Z. Y., Knorr, D. A., Bendzick, L., Allred, J. & Kaufman, D. S. (2014). Expression of Chimeric Receptor CD4 zeta by Natural Killer Cells Derived from Human Pluripotent Stem Cells Improves In Vitro Activity but Does Not Enhance Suppression of HIV Infection In Vivo. Stem cells 32, 2014.

16. Sugimura, R., Jha, D. K., Han, A., Soria-Valles, C., da Rocha, E. L., Lu, Y. F., Goettel, J. A., Serrao, E., Rowe, R. G., Malleshaiah, M., Wong, I., Sousa, P., Zhu, T. N., Ditadi, A., Keller, G., Engelman, A. N., Snapper, S. B., Doulatov, S. & Daley, G. Q. (2017). Haematopoietic stem and progenitor cells from human pluripotent stem cells. Nature 545, 2017.

17. Woll, P. S., Martin, C. H., Miller, J. S. & Kaufman, D. S. (2005). Human embryonic stem cell-derived NK cells acquire functional receptors and cytolytic activity. Journal of immunology 175, 2005.

18. Hermanson, D. L., Ni, Z. & Kaufman, D. S. (2015). Human Pluripotent Stem Cells as a Renewable Source of Natural Killer Cells. In Hematopoietic Differentiation of Human Pluripotent Stem Cells (Cheng, T., ed), pp. 69–79, Springer Netherlands, Dordrecht.

19. Hermanson, D. L., Bendzick, L., Pribyl, L., McCullar, V., Vogel, R. I., Miller, J. S., Geller, M. A. & Kaufman, D. S. (2016). Induced Pluripotent Stem Cell-Derived Natural Killer Cells for Treatment of Ovarian Cancer. Stem Cells 34, 2016.

20. Denman, C. J., Senyukov, V. V., Somanchi, S. S., Phatarpekar, P. V., Kopp, L. M., Johnson, J. L., Singh, H., Hurton, L., Maiti, S. N., Huls, M. H., Champlin, R. E., Cooper, L. J. & Lee, D. A. (2012). Membrane-bound IL-21 promotes sustained ex vivo proliferation of human natural killer cells. PLoS One 7, 2012.

21. Ng, E. S., Davis, R., Stanley, E. G. & Elefanty, A. G. (2008). A protocol describing the use of a recombinant protein-based, animal product-free medium (APEL) for human embryonic stem cell differentiation as spin embryoid bodies. Nat Protoc 3, 2008.

22. Claassen, D. A., Desler, M. M. & Rizzino, A. (2009). ROCK inhibition enhances the recovery and growth of cryopreserved human embryonic stem cells and human induced pluripotent stem cells. Molecular reproduction and development 76, 2009.

23. Zou, L., Chen, Q. S., Quanbeck, Z., Bechtold, J. E. & Kaufman, D. S. (2016). Angiogenic activity mediates bone repair from human pluripotent stem cell-derived osteogenic cells. Sci Rep-Uk 6, 2016.

24. Ludwig, T. E., Bergendahl, V., Levenstein, M. E., Yu, J., Probasco, M. D. & Thomson, J. A. (2006). Feeder-independent culture of human embryonic stem cells. Nature methods 3, 2006.

25. Hexum, M. K., Tian, X. & Kaufman, D. S. (2011). In vivo evaluation of putative hematopoietic stem cells derived from human pluripotent stem cells. Methods in molecular biology 767, 2011.

26. Watanabe, K., Ueno, M., Kamiya, D., Nishiyama, A., Matsumura, M., Wataya, T., Takahashi, J. B., Nishikawa, S., Nishikawa, S., Muguruma, K. & Sasai, Y. (2007). A ROCK inhibitor permits survival of dissociated human embryonic stem cells. Nature biotechnology 25, 2007.

